# Dual role of *Toxoplasma gondii* ROP5 and ROP18 for NLRP3 inhibition

**DOI:** 10.1101/2023.09.17.558105

**Authors:** Mateo Murillo Léon, Aura María Bastidas Quintero, Shishir Singh, Xochitl Müller, Oliver Gorka, Benedikt Simon Saller, Ailan Farid Arenas Soto, Claudia Campos, Jorge Enrique Gomez-Marin, Olaf Groß, Klaus Pfeffer, Jonathan Charles Howard, Gregory Alan Taylor, Tobias Steinfeldt

**Affiliations:** Institute of Virology, Medical Center University of Freiburg, 79104 Freiburg, Germany; Faculty of Medicine, University of Freiburg, 79104 Freiburg, Germany; Faculty of Biology, University of Freiburg, 79104 Freiburg, Germany; Henry D. Janowitz Division of Gastroenterology, Department of Medicine, Icahn School of Medicine at Mount Sinai, New York, NY, United States of America; Institute of Neuropathology, Medical Center-University of Freiburg, Faculty of Medicine, University of Freiburg, 79104 Freiburg, Germany; Grupo GEPAMOL, Centro de Investigaciones Biomedicas, Universidad del Quindio, Armenia, Quindio, Colombia; Fundacao Calouste Gulbekian, Instituto Gulbekian de Ciencia, 2780-156 Oeiras, Portugal; Signalling Research Centers BIOSS and CIBSS, University of Freiburg, 79104 Freiburg, Germany; Institute of Medical Microbiology and Hospital Hygiene, Heinrich-Heine-University Düsseldorf, 40225 Düsseldorf, Germany; Departments of Medicine; Molecular Genetics and Microbiology; and Immunology; and Center for the Study of Aging and Human Development, Duke University Medical Center, NC 27705 Durham, North Carolina, United States of America; Geriatric Research, Education, and Clinical Center, Durham VA Health Care System, NC 27705 Durham, North Carolina, United States of America

**Keywords:** NLRP3, *Toxoplasma gondii*, inflammasome activation, inflammasome inhibition, host cell resistance molecule GBP5, parasite virulence effectors ROP5/ROP18

## Abstract

Inflammasome activation leads release of IL-1β, a proinflammatory cytokine that drives antimicrobial immune responses. *Toxoplasma gondii* has been shown to activate the NLRP3 inflammasome but the trigger has not yet been identified. Here we provide evidence that vacuolar disruption is a prerequisite for NLRP3 activation. *T*. *gondii* type I ROP5 and ROP18 protect the parasitophorous vacuolar membrane (PVM) and thereby inhibit inflammasome activation and IL-1β release. Besides protection of the PVM, we demonstrate an additional function of ROP5 and ROP18 for NLRP3 inhibition. We demonstrate the molecular mechanism of this inhibition includes direct interaction with GBP5. In conclusion, *T. gondii* ROP5 and ROP18 inhibit IL-1β production by protection of the intracellular replicative niche of the parasite and by actively subverting NLRP3 activation. Our findings provide further insight into the intricate mechanisms governing inflammasome activation and inhibition, enhancing our understanding of the complex dynamics during *T*. *gondii* infection.

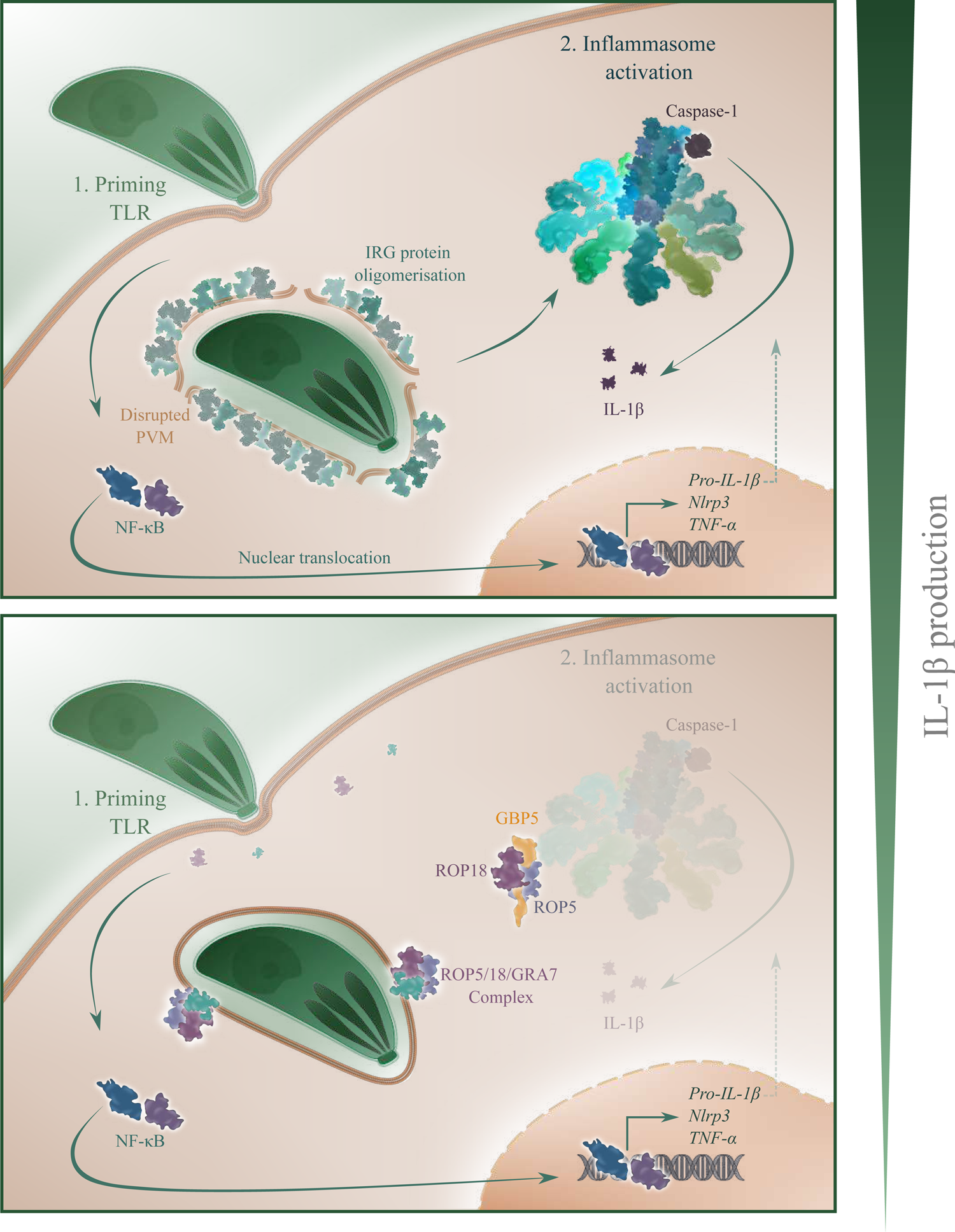

## Introduction

Interleukin-1 beta (IL-1β) is a pro-inflammmatory cytokine that drives antimicrobial immune responses. Dysregulation of pro-inflammatory cytokine production, such as IL-1β, can cause tissue damage and severe immunopathology and must therefore be tightly regulated. To this end, a nuclear factor kappa B (NF-κB)-dependent priming signal induces pro-IL-1β production. A second signal triggers its proteolytic processing to bioactive IL-1β by cytoplasmic complexes called inflammasomes. Inflammasomes are part of the innate immune system and play a pivotal role in response to infection and perturbation of cellular homeostasis. A minimal inflammasome is usually composed of an upstream sensor belonging either to the NLR (nucleotide-binding oligomerisation domain (NOD)-like receptor), NAIP (neuronal-apoptosis inhibitor protein) or ALR (AIM2-like receptor) families, the adaptor protein ASC (apoptosis-associated speck-like proteins containing CARDs (caspase recruitment domains)), and an effector caspase. In response to microbial infection or tissue damage, danger- or pathogen-associated molecular patterns (DAMPs or PAMPs) lead to TLR (toll-like receptor) activation and NF-κB signaling resulting in transcription of pro-IL-1β and pro-IL-18. Once activated by DAMPs or PAMPs, inflammasomes start to assemble into large multiprotein complexes that serve as platforms for autoproteolysis and thereby activation of the cysteine protease pro-caspase-1. Caspase-1 activation results in processing of IL-1β and IL-18 along with gasdermin-D (GSDMD) and eventually initiation of pyroptosis, a proinflammatory cell death pathway [1].

Interferon (IFN)-γ-inducible Guanylate Binding Proteins (GBP proteins) and Immunity-Related GTPases (IRG proteins) were identified as regulators of canonical and non-canonical inflammasomes following infection by pathogenic bacteria [2–5] or *Toxoplasma gondii* (*T. gondii*) [6]. In this regard, human and mouse GBP5 (these orthologs share 67 % identity at the amino acid level [7]) associates directly with NLRP3 (nucleotide-binding oligomerization domain, leucine rich repeat and pyrin domain containing 3) upon infection with *Salmonella typhimurium* or *Listeria monocytogenes*, promoting ASC oligomerisation and thereby full NLRP3 inflammasome activation [8]. Different expression levels of strain-specific *Gbp* genes [9, 10] adjacent to *Gbp5* might be an explanation why this finding could not be reproduced in other studies [11, 12]. The contribution of GBP5 to IL-1β release upon *T. gondii* infection was independent of IFN-γ and classical inflammasome activation [6], yet its molecular function remains elusive.

*T. gondii* is an obligate intracellular food-borne parasite with a broad host range that can cause severe disease, particularly in immune compromised individuals. On a global scale, one third of the human population as well as many wild mammals and birds are infected with *T*. *gondii*. Of the three clonal lineages that are commonly found in Europe and North America, type I strains are highly virulent for laboratory mice at low inocula (LD_100_=1) whereas type II and III strains are less virulent (LD_50_=10^3^-10^5^). Control of *T. gondii* infection in mice requires production of the pro-inflammatory cytokine IL-12 following recognition of parasite-derived profilin-like protein and cyclophilin-18 by TLR11 and TLR12 in the acute phase of infection. IL-12 production leads to induction of IFN-γ, a key mediator of immunity to *T. gondii*. In genetic crosses between different *T. gondii* clonal lineages, the serine-threonine kinase ROP18 and a family of highly polymorphic ROP5 pseudokinases emerged as major virulence effectors in laboratory mice [13–15]. These and other parasite-derived effector proteins [16–28] prevent the accumulation of IRG and GBP proteins at the parasitophorous vacuolar membrane (PVM) that otherwise induce vacuole and parasite membrane disintegration, inevitably followed by acute death of the parasite and host cell death [29–32]. The crucial role of IRG and GBP proteins in this regard is supported by the fact that in the absence of IRG protein accumulation at the PVM, membrane rupture and host cell death was never observed [29].

In human cells, resistance against *T. gondii* is mediated by depletion of tryptophan [33, 34]. Furthermore, GBP1, GBP2 and GBP5 have been demonstrated to control infection in human macrophages [32, 35].

*T. gondii* is an activator of the NLRP1 and NLRP3 inflammasomes *in vivo* [36, 37] and an association of polymorphisms in human *Nlrp1* and *P2RX7* with susceptibility to congenital and ocular toxoplasmosis was reported [38–43]. Modulation of inflammasomes is critical to promote pathogen virulence and like many other vertebrate adapted pathogens, *T. gondii* inhibits IL-1β production. *T. gondii* type I strain infections result in reduced NF-κB signaling and NLRP3 activation in human neutrophils [44] and *T. gondii* type I GRA16 was identified as inhibitor of NF-κB signaling in human NCI-H1299 cells [45]. Furthermore, GRA15 contributes to over-induction of pro IL-1β by type II but not type I and type III strains in human monocytes [46]. These studies emphasize an important role of IL-1β for *T. gondii* control and document differences between canonical parasite strains. In contrast, no differences between canonical *T. gondii* lineages with respect to IL-1β release have been observed in bone marrow-derived macrophages (BMDMs) [6, 37], even in the presence of IFN-γ [47]. Furthermore, the molecular trigger that initiates inflammasome activation remains elusive and the contribution of GBP5 to IL-1β release [6] during murine and human *T. gondii* infections has not yet been further investigated.

Here we demonstrate that vacuolar disruption is a prerequisite for NLRP3 inflammasome activation upon *T. gondii* infection in bone marrow-derived dendritic cells (BMDCs). *T. gondii* type I effectors ROP5 and ROP18 protect the PVM from IRG/GBP-mediated destruction explaining differences in inflammasome activation and consequently IL-1β secretion between *T. gondii* wt and *rop5* or *rop18* ko strains. In addition to protection of the PVM, ROP5 and ROP18 can directly inhibit NLRP3 inflammasome assembly and in case of ROP5, this direct inhibition is mediated via binding to GBP5. These results demonstrate a dual role of ROP5/ROP18 for inflammasome inhibition in BMDCs. We further demonstrate the requirement of GBP5 for inflammasome activation in *T. gondii* infection of human macrophages. However, in human cells the molecular function of GBP5 is not inhibited by ROP5 or ROP18. Our findings deepen the understanding of the complex relationship between the host’s immune response and the virulence strategies employed by *T. gondii*.

## Results

### *T. gondii* type I effectors ROP5 and ROP18 inhibit IL-1β secretion

It has been demonstrated that mouse macrophages secrete IL-1β in response to all three clonal lineages of *Toxoplasma gondii* (*T. gondii*) but apparent differences between strains could not be observed [6, 37, 47]. In mice, type I *Toxoplasma gondii* (*T. gondii*) effectors ROP5 and ROP18 protect the parasitophorous vacuolar membrane (PVM) against disruption by interferon (IFN)-γ-inducible Guanylate Binding Proteins (GBP proteins) and Immunity-Related GTPases (IRG proteins). Therefore, IL-1β release in presence and absence of ROP5 or ROP18 should be informative whether PVM disruption is a necessary trigger for inflammasome activation upon *T. gondii* infection. Bone marrow-derived dendritic cells (BMDCs) prepared from C57BL/6 mice were seeded in dishes of a 96 well-plate in the absence or presence of IFN-γ. After stimulation with 20 ng/ml LPS for 3 h to achieve saturated pro-IL-1β levels, cells were infected with *T. gondii* RHΔ*hxgprt*, RHΔ*rop5* or RHΔ*rop18* for 6 h and IL-1β levels determined in the supernatants by quantitative ELISA (Fig. 1A, see also Suppl. Fig. 1 for absolute IL-1β levels) and Western blot (Fig. 1B and Suppl. Fig. 2). In the absence of IFN-γ, IL-1β levels are significantly increased in RHΔ*rop5* compared with wt strain-infected cells. IFN-γ-treatment induced an overall increase of IL-1β secretion and differences between *T. gondii* wt and ko strains were even more pronounced (Fig. 1A, 1B and Suppl. Fig. 2). To investigate whether elevated IL-1β levels in the supernatants are due to inflammasome activation, mature caspase-1 release was determined upon infection with the same *T. gondii* strains. Western blot analysis reveals elevated levels of mature caspase-1 in the supernatants upon *T. gondii* ko compared to wt strain infections (Fig. 1B and Suppl. Fig. 2). Differences between the wild-type (wt) and knockout (ko) *T. gondii* strains are not due to differences in infection rates (Fig. 1B, GRA7 protein levels) or differences in nuclear factor kappa B (NF-κB) activity determined by tumor necrosis factor-alpha (TNF-α) secretion (Fig. 1C). IL-1β levels in the presence and absence of IFN-γ directly correlate with percentage of cell death - representative for vacuolar disruption - upon *T. gondii* infection measured by lactate dehydrogenase (LDH) release (Fig. 1D). To determine the impact of different inflammasome components on increased IL-1β levels in case of *T. gondii* RHΔ*rop5* or RHΔ*rop18*, IL-1β levels were measured in the supernatants of infected BMDCs deficient for either ASC, NLRP3 or caspase-1. IL-1β levels in the supernatant of all infected ko cells are significantly reduced compared to wt cells indicating that NLRP3 contributes to the inflammatory response upon *T. gondii* infection (Fig. 2A). Differences between the wt and ko *T. gondii* strains are not due to differences in NF-κB activity (Fig. 2B). *T. gondii* ko strain infections in the presence of IFN-γ lead to increased levels of cleaved gasdermin-D (GSDMD) in comparison to the wt strain (Suppl. Fig. 2) implicating increased levels of GSDMD-mediated pyroptosis [48]. However, the absence of inflammasome components does not result in diminished cell death upon wt and ko strain infections compared to wt cells (Fig. 2C). Furthermore, we observed no discernible differences between wt and *GSDMD* ko cells concerning cell death following infection with the corresponding *T. gondii* strains (Suppl. Fig. 3). These results are congruent with pyroptosis-independent IL-1β secretion and further support the findings that IRG/GBP-mediated PVM disruption is the cause of cell death in our experiments.

**Figure 1.**
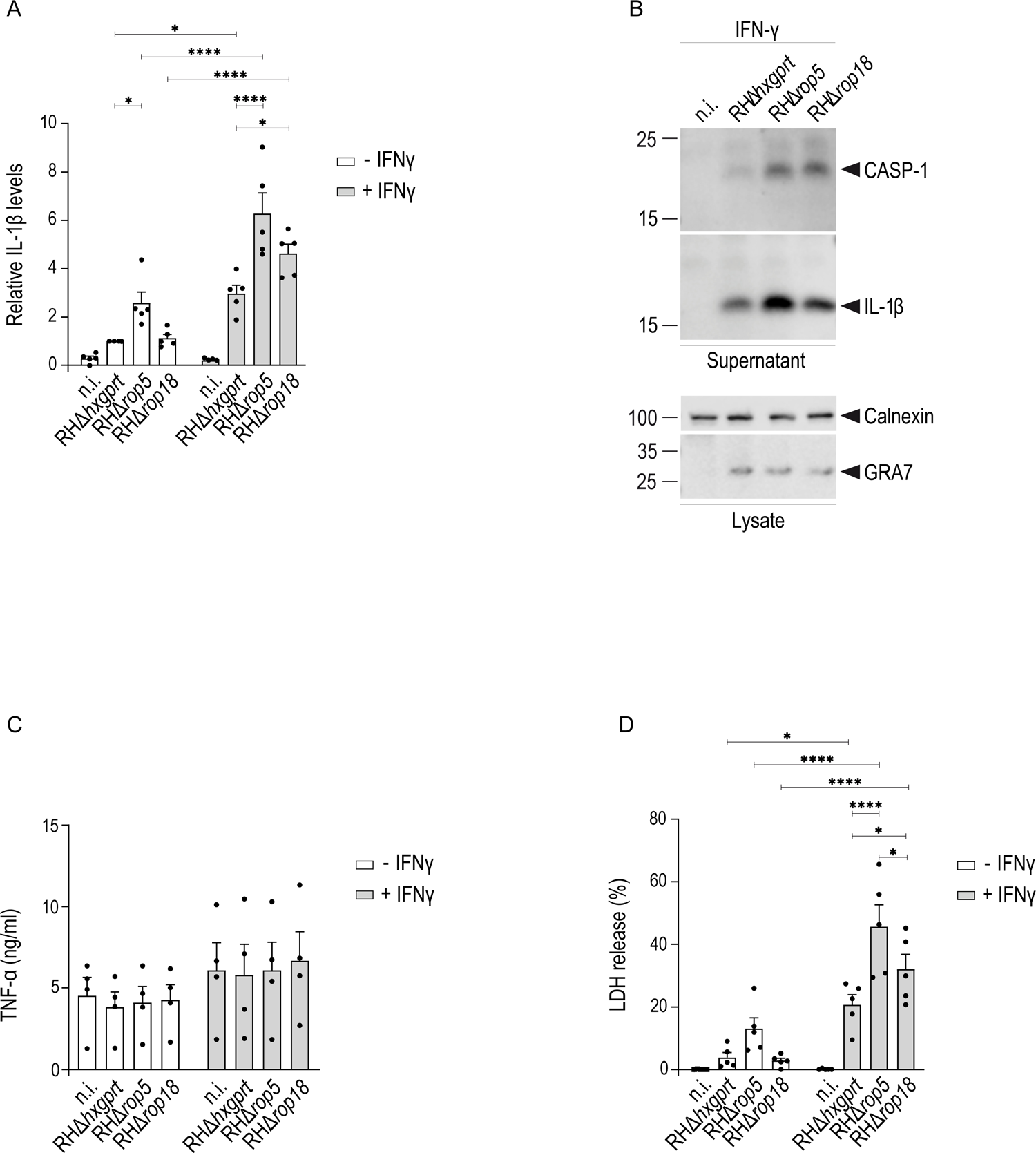
*T. gondii* ROP5 and ROP18 inhibit IL-1β production. BL/6 BMDCs were primed for 3 h with LPS and subsequently infected with indicated *T. gondii* strains in presence or absence of IFN-γ for 6 h. A, B, IL-1β levels in the supernatants are significantly increased upon *T. gondii* infection in the presence of IFN-γ compared with unstimulated conditions (A). Comparing *T. gondii* strains, IL-1β secretion is significantly increased in ko compared to wt strain infections in the presence of IFN-γ and in *T. gondii* RHΔ*rop5* versus the wt strain in the absence of IFN-γ. Non-infected cells served as negative control for inflammasome activation. C, No differences in TNF-α secretion are detectable between *T. gondii* strains. D, Cell death, determined by LDH release into the supernatant, is significantly increased in *T. gondii* RHΔ*rop5* and RHΔ*rop18* compared with wt strain infections in the presence but not in the absence of IFN-γ. A, C, D, Error bars indicate the mean and standard error of the mean (SEM) of five (A, D) or four (C) independent experiments performed in triplicates. One-way analysis of variance (ANOVA) followed by Tukey’s (A) or Holm-Sidak’s multiple comparison (D) was used to test differences between groups; ****p < 0.0001; *p < 0.05.

**Figure 2.**
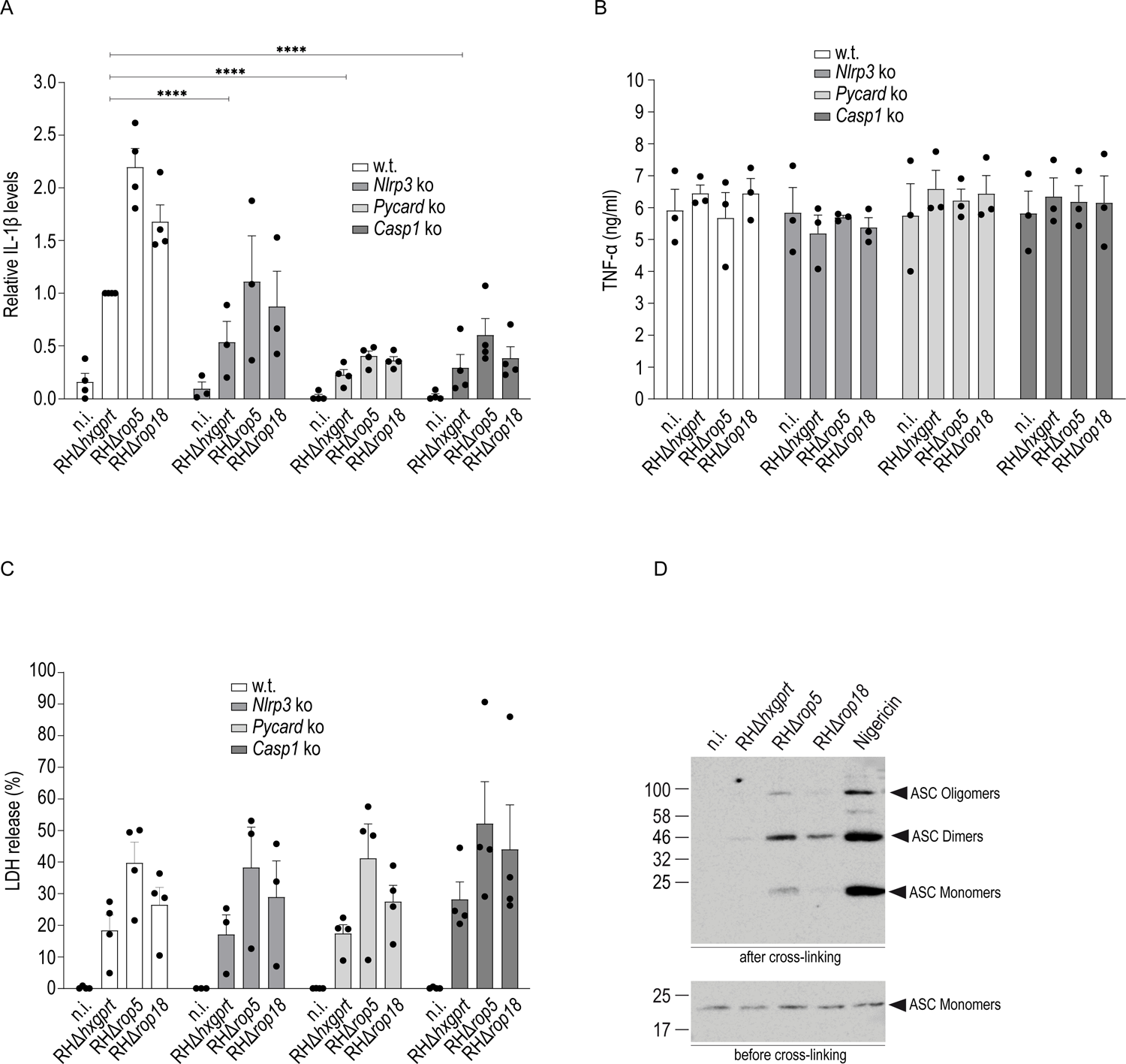
*T. gondii* ROP5 and ROP18 inhibit the NLRP3 inflammasome. BL/6 BMDCs were prestimulated for 3 h with LPS and subsequently infected with indicated *T. gondii* strains in presence of IFN-γ for 6 h. A, Compared to wt cells, IL-1β levels in the supernatants are significantly decreased in the absence of NLRP3 (*Nlrp3* ko), ASC (*Pycard* ko) or caspase-1 (*Casp1* ko) upon infection with *T. gondii* wt, RHΔ*rop5* or RHΔ*rop18*. B, No differences in TNF-α secretion are detectable between wt and ko cell lines upon infection with *T. gondii* wt, RHΔ*rop5* or RHΔ*rop18*. C, No differences in LDH release into the supernatant are detectable between wt and ko cell lines upon infection with *T. gondii* wt, RHΔ*rop5* or RHΔ*rop18*. Error bars indicate the mean and standard error of the mean (SEM) of four independent experiments. One-way analysis of variance (ANOVA) followed by Tukey’s multiple comparison was used to test differences between groups; ****p < 0.0001; ***p < 0.001; **p < 0.01. D, ASC multimerisation is increased upon RHΔ*rop5* or RHΔ*rop18* compared with wt strain infections. Nigericin treatment served as positive control for inflammasome activation. In all cases, non-infected cells served as negative control for inflammasome activation.

The adaptor ASC is crucial for canonical inflammasome activation by mediating the interaction between NLRP3 and caspase-1. Upon activation, NLRP3 nucleates ASC to form particular foci leading to cleavage of caspase-1. To test whether ASC oligomerization and inflammasome assembly is disturbed upon *T. gondii* infection, BMDCs seeded in presence of IFN-γ and LPS were infected with *T. gondii* wt RHΔ*hxgprt*, RHΔ*rop5* or RHΔ*rop18* strains. At 3 h post infection, cell lysates were prepared and ASC cross-linking performed with 2 mM DSS for 30 min at RT. ASC oligomerization was analysed by Western blot using an ASC-specific antibody. ASC dimer, tetramer and higher order oligomer formation is reduced in case of RHΔ*hxgprt* compared to RHΔ*rop5* or RHΔ*rop18* infections (Fig. 2D). Nigericin served as positive control for ASC oligomerization.

In summary, these results demonstrate that NLRP3-mediated IL-1β production upon *T. gondii* infection can efficiently be inhibited in the presence of type I parasite effectors ROP5 and ROP18 that protect the PVM from IRG/GBP-mediated disruption. IL-1β levels in the absence of *Nlrp3* confirms the activation of other inflammasomes upon *T. gondii* infection [37, 47, 49].

### *T. gondii* PVM disruption is a prerequisite for activation of the NLRP3 inflammasome

IFN-γ triggers expression of IRG and GBP proteins in moue cells that contribute to disruption of the PVM and subsequent parasite and host cell death [29]. Increased IL-1β release in the presence of IFN-γ (Fig. 1) suggests that PVM disruption is an important trigger for inflammasome activation upon *T. gondii* infection. Regulator IRG proteins Irgm1, Irgm2 and Irgm3 keep IRG effectors, that accumulate at the *T. gondii* PVM, in the inactive GDP-bound state at endomembranes in non-infected cells. In the absence of IRG regulators, effector IRG and GBP proteins do not accumulate at the PVM and membrane integrity is preserved [50–54]. To investigate whether PVM disruption is a prerequisite for inflammasome activation, IL-1β release was determined upon infection of *Irgm1*/*Irgm3* ko BMDCs. Under these conditions, a significant decrease in IL-1β release (Fig. 3A) and cell death (Fig. 3B) can be detected upon infection but not upon nigericin treatment (Fig. 3C). Furthermore, no significant differences between *T. gondii* wt and ko strains are apparent, consistent with PVM disruption contributing to inflammasome activation upon *T. gondii* infection of BMDCs.

**Figure 3.**
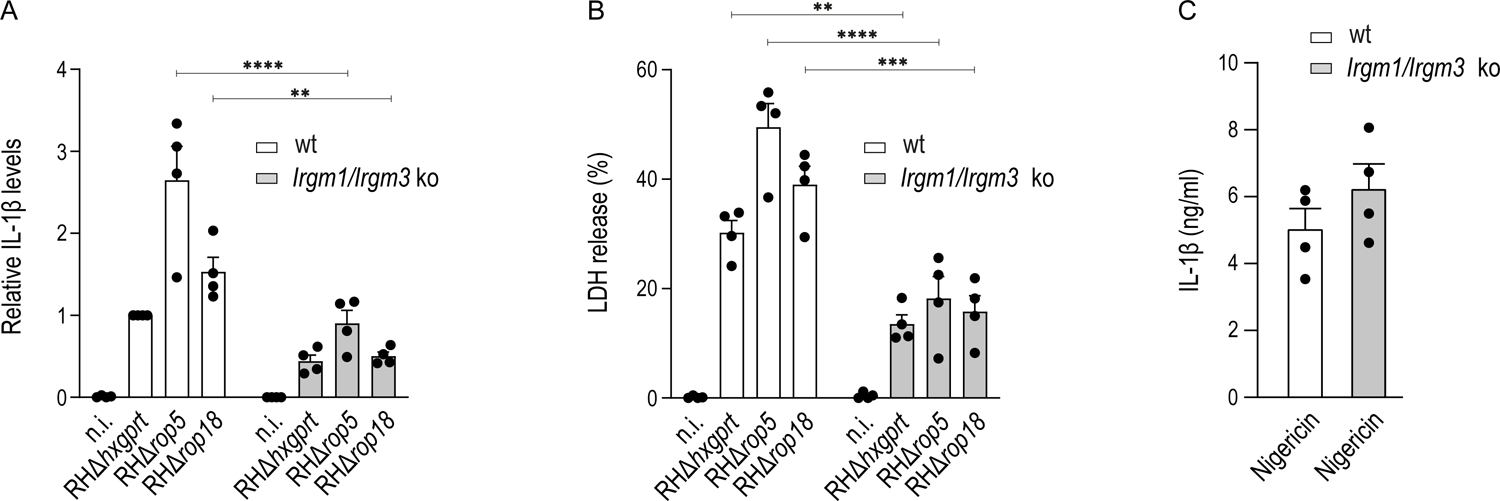
*T. gondii* PVM disruption is a prerequisite for activation of the NLRP3 inflammasome. BL/6 wt or *Irgm1/Irgm3* ko BMDCs were prestimulated for 3 h with LPS and subsequently infected with indicated *T. gondii* strains in presence of IFN-γ for 6 h. Compared to wt cells, IL-1β (A) or LDH release (B) into the supernatant is significantly decreased in the absence of Irgm1/Irgm3 upon infection with *T. gondii* wt, RHΔ*rop5* or RHΔ*rop18*. Non-infected cells served as negative control for inflammasome activation. Error bars indicate the mean and standard error of the mean (SEM) of four independent experiments. One-way analysis of variance (ANOVA) followed by Tukey’s multiple comparison was used to test differences between groups; ****p < 0.0001; ***p < 0.001; **p < 0.01. C, No differences in IL-1β levels between wt and *Irgm1/Irgm3* ko cells are apparent in upon nigericin treatment.

### IFN-α contributes to IRG-mediated PVM disruption and inflammasome activation

We have shown here that *T. gondii* PVM disruption mediated by vacuolar IRG/GBP protein accumulation upon IFN-γ induction triggers inflammasome activation (Fig. 3). Nevertheless, RHΔ*rop5* infections result in increased IL-1β and LDH levels compared with RHΔ*hxgprt* even in the absence of IFN-γ (Fig. 1A). However, LPS induction or *T. gondii* infection under these conditions may also induce IFN-α [55]. To determine the contribution of IFN-α to inflammasome activation, we treated BMDCs freshly prepared from C57BL/6 wt and IFN-α receptor (*Ifnar*) ko mice with LPS for 3 h and determined IL-1β release into the supernatant 6 h post *T. gondii* infection. In wt cells, significantly higher IL-1β levels in the supernatants can be detected upon RHΔ*rop5* infection compared with RHΔ*hxgprt* whereas no differences between strains are apparent in *Ifnar* ko cells (Fig. 4A). Nigericin treatment served as positive control for inflammasome activation. As expected, IL-1β release upon nigericin-treatment is reduced in wt cells compared with *Ifnar* ko cells (Fig. 4B) because type I IFN has been described as inhibitor of inflammasome activation [56]. IL-1β levels directly correlate with percentage of cell death upon *T. gondii* infection measured by LDH release (Fig. 4C). To confirm that IL-1β release under these conditions is due to inflammasome activation mediated by IRG/GBP protein-dependent PVM disruption, we determined IRG protein expression levels by Western blot and PVM accumulation by immunofluorescence in *Ifnar* ko and wt cells upon IFN-α-treatment. IFN-α induces the expression of Irga6 and Irgb6 in wt but not *Ifnar* ko cells (Fig. 4D) and Irga6-positive RHΔ*rop5*-derived vacuoles can be detected in wt BMDCs (Fig. 4E). These results demonstrate that IL-1β release upon *T. gondii* infection in the absence of IFN-γ (Fig. 1) is due to IFN-α-inducible IRG/GBP proteins that accumulate at the PVM and offer an explanation for IL-1β release in previous studies [6, 37, 47].

**Figure 4.**
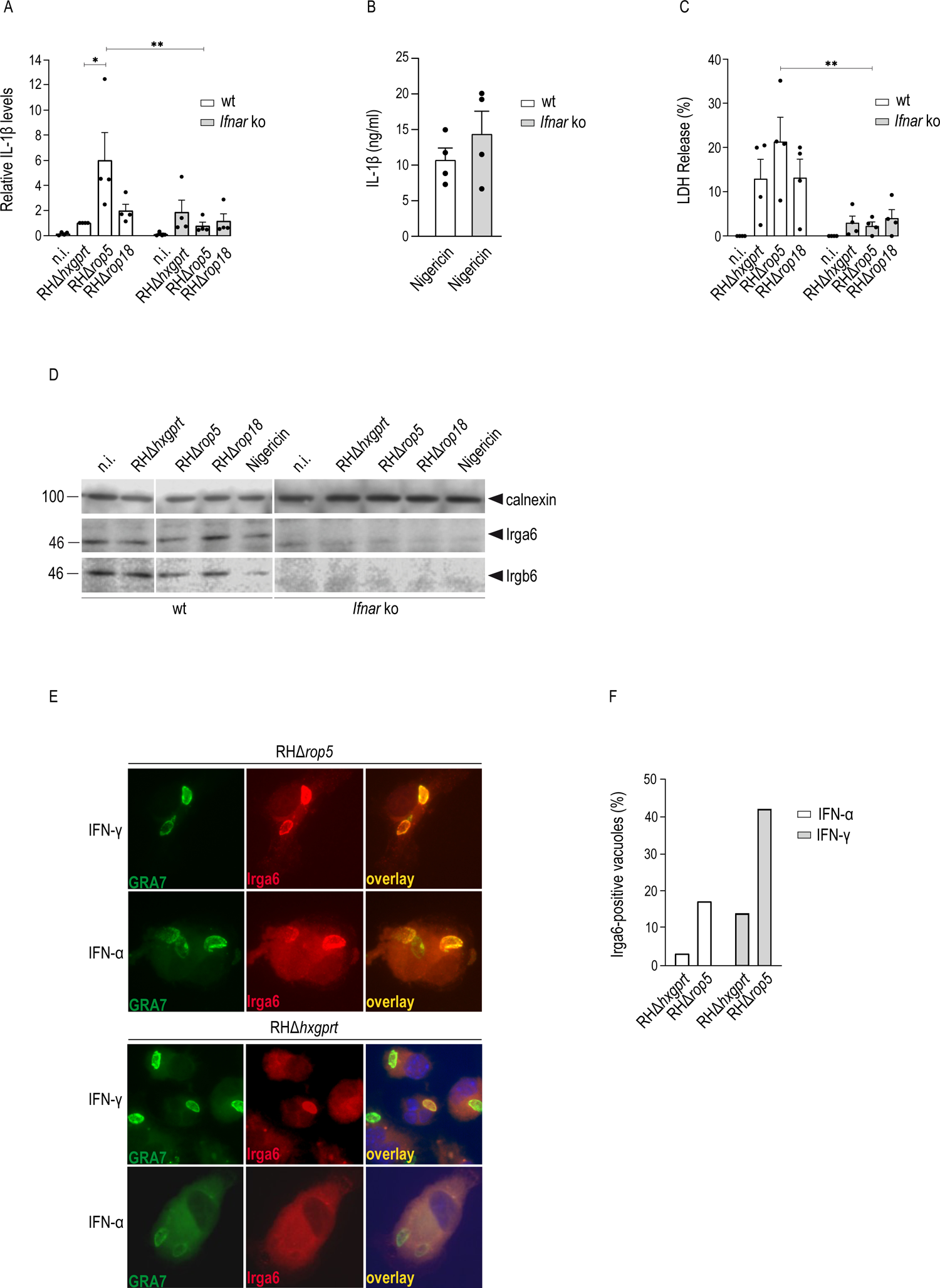
IFN-α contributes to IRG-mediated PVM disruption and inflammasome activation. BL/6 wt or *Ifnar* ko BMDCs were prestimulated with LPS and IFN-α and subsequently infected with indicated *T. gondii* strains for 6 h. A, IL-1β levels in the supernatants are significantly increased in wt cells upon *T. gondii* RHΔ*rop5* infection compared with wt and RHΔ*rop18* strains. In the absence of the IFN-α receptor, no differences of IL-1β levels between wt and RHΔ*rop5* parasite strains are detectable anymore. Non-infected cells served as negative control for inflammasome activation. B, Nigericin treatment served as positive control for inflammasome activation. C, Differences in LDH release into the supernatant between *T. gondii* wt and RHΔ*rop5* strains in wt cells are equalised upon *T. gondii* infection of *Ifnar* ko cells. A, C, Error bars indicate the mean and standard error of the mean (SEM) of four independent experiments. One-way analysis of variance (ANOVA) followed by Tukey’s multiple comparison was used to test differences between groups; **p < 0.01; *p < 0.05. D, Expression of IRG effectors Irga6 and Irgb6 can be detected by Western blot with specific antibodies in wt but not *Ifnar* ko cells upon IFN-α treatment. E, F, Irga6-positive vacuoles in BMDCs. BMDCs were stimulated with IFN-α or IFN-γ for 24 h before infection with RHΔ*hxgprt* or RHΔ*rop5* at an MOI of 3 for 2 h. Representative fluorescent images of RHΔ*hxgprt*- and RHΔ*rop5*-derived vacuoles. GRA7 in green, Irga6 in red and overlay in yellow (E). Frequencies of Irga6-positive RHΔ*hxgprt*- and RHΔ*rop5*-derived vacuoles (F).

### *T. gondii* ROP5 and ROP18 inhibit the NLRP3 inflammasome

We have demonstrated previously that Irgb2-b1, one of the most polymorphic IRG family members identified so far, is largely responsible for control of *T. gondii* type I strain infections in CIM mice, a wild-derived *Mus musculus castaneus* subspecies [57]. Irgb2-b1 counters parasite virulence by directly binding to ROP5 isoform B (ROP5B). Consequently, effector IRG proteins can accumulate at the PVM leading to its disintegration and parasite and host cell death [57, 58]. With that premise, the level of inflammasome activation triggered by PVM disruption should be indistinguishable between *T. gondii* type I wt and RHΔ*rop5* or RHΔ*rop18* strain infections in BMDCs derived from CIM mice. We infected BMDCs from C57BL/6 and CIM mice seeded in absence or presence of IFN-γ and after LPS treatment with *T. gondii* RHΔ*hxgprt*, RHΔ*rop5* or RHΔ*rop18* for 6 h and determined IL-1β and LDH release into the supernatants. The level of IL-1β release (Fig. 5A) and cell death (Fig. 5B) is significantly increased in case of RHΔ*rop5* or RHΔ*rop18* compared with wt strain infected C57BL/6 cells, confirming our previous findings. In CIM cells, cell death upon wt and ko strain infections is essentially the same, confirming the CIM-inherent resistance mechanism against virulent *T. gondii* strains (Fig. 5B). Interestingly, however, IL-1β levels still remain significantly higher in RHΔ*rop5* or RHΔ*rop18* infected CIM BMDCs compared with wt strain infections (Fig. 5A).

**Figure 5.**
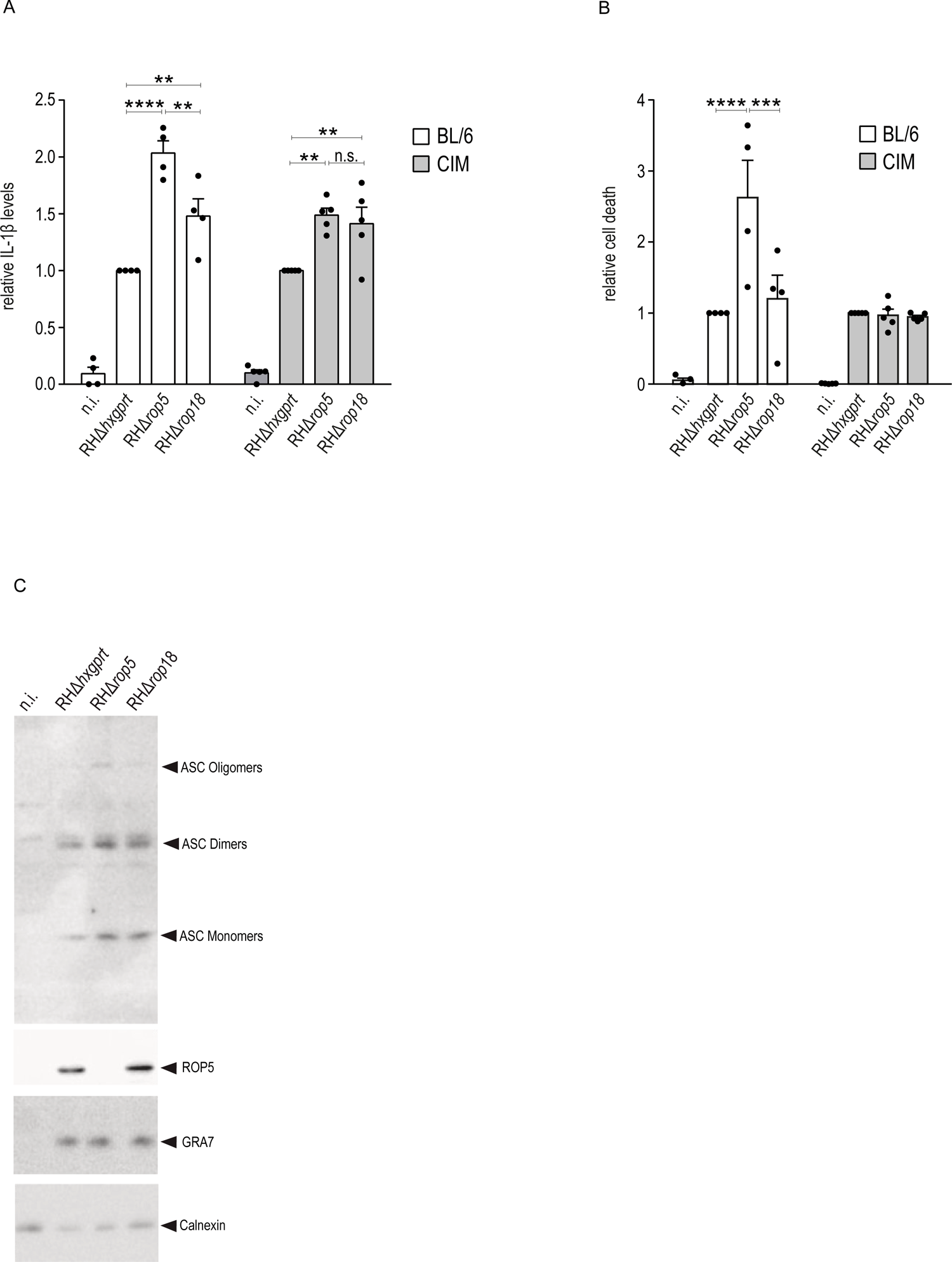
*T. gondii* ROP5 and ROP18 inhibit the NLRP3 inflammasome. BL/6 or CIM BMDCs were prestimulated for 3 h with LPS and subsequently infected with indicated *T. gondii* strains in presence of IFN-γ for 6 h. A, Compared to BL/6 cells, levels of LDH in the supernatant are indistinguishable between *T. gondii* strains in CIM cells. B, IL-1β levels in the supernatants are significantly increased in case of *T. gondii* RHΔ*rop5* and RHΔ*rop18* compared with wt strain infections in BL/6 and CIM cells. A, B, Error bars indicate the mean and standard error of the mean (SEM) of four or five independent experiments. One-way analysis of variance (ANOVA) followed by Tukey’s multiple comparison was used to test differences between groups; ****p < 0.0001; **p < 0.01. C, ASC multimerisation is increased upon RHΔ*rop5* and RHΔ*rop18* compared with wt strain infections in CIM cells. In all cases, non-infected cells served as negative control for inflammasome activation.

Similar to C57BL/6 BMDCs (Fig. 2D), ASC oligomerisation is inhibited in RHΔ*hxgprt* but not RHΔ*rop5* or RHΔ*rop18* infections of CIM BMDCs (Fig. 5C). These results imply that inflammasome activation during *T. gondii* infection may be triggered by factors other than PVM disruption. Alternatively, there might be an additional inhibitory role of ROP5/ROP18 in inflammasome activation. This inhibitory function appears to be distinct from ROP5/ROP18-mediated inactivation of IRG/GBP proteins, which serves to protect against PVM disruption.

### GBP5 interacts directly with ROP5 and contributes to inflammasome activation upon *T. gondii* infection in BMDCs

In an unbiased approach, we screened all human and murine GBP proteins (hGBP and mGBP) in combination with *T. gondii* ROP5 and identified direct interaction of ROP5A and ROP5B with hGBP5 and mGBP5 (data not shown). Mouse and human GBP5 have been demonstrated to mediate selective inflammasome assembly and activation in response to pathogenic intracellular bacteria [7]. To determine a possible influence of GBP5 on inflammasome activation upon *T. gondii* infection and an inhibitory function of ROP5 and ROP18 in this context, we reinvestigated murine GBP5 (mGBP5) and human GBP5 (hGBP5) for direct interaction with these parasite effectors. In a Yeast Two-Hybrid (YTH) approach, ROP5A and ROP5B are shown to interact with mGBP5 and hGBP5 (Fig. 6A and 6B). In comparison to mGBP5, the ROP5:hGBP5 interaction was rather weak and was therefore confirmed in pull-down experiments with a GST-tagged hGBP5 fusion protein on *T. gondii* type I detergent lysates and subsequent Western blot analysis using a specific anti-ROP5 antibody (Fig. 6C).

**Figure 6.**
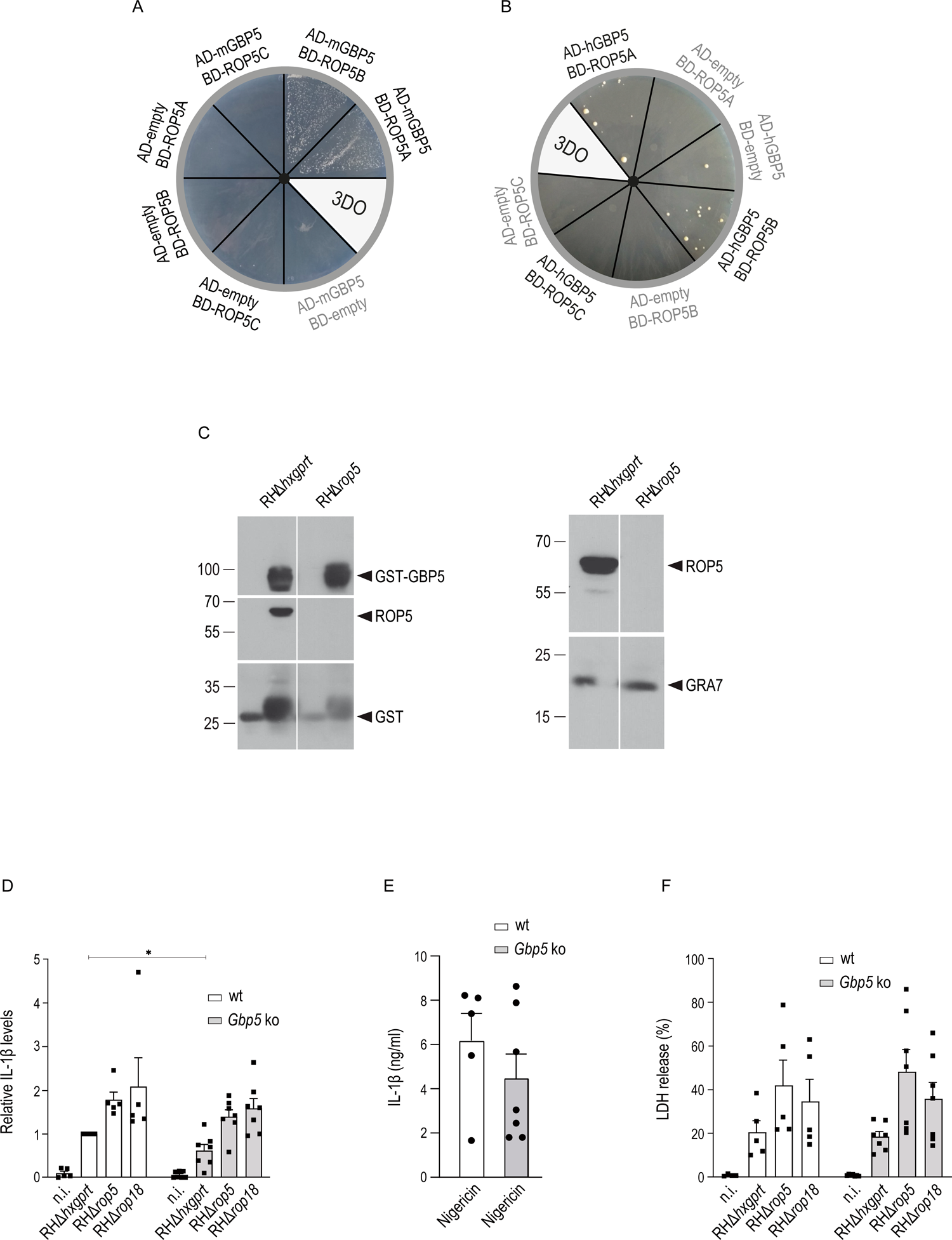
GBP5 interacts directly with ROP5 and contributes to inflammasome activation upon *T. gondii* infection in BMDCs. A, B, Yeast Two-Hybrid. *T. gondii* ROP5A and ROP5B directly bind to mGBP5 (A) and hGBP5 (B). Proteins were expressed either as fusion to the Gal4 transcriptional activation (AD) or DNA-binding (BD) domain. Colony growth under 3DO conditions demonstrates protein:protein-interaction. C, Pull-down of ROP5 by GST-hGBP5 from RH tachyzoite detergent lysates. RHΔ*rop5* tachyzoite detergent lysates were included as control (left-hand panel). Similar ROP5 and GRA7 expression levels in tachyzoite detergent lysates used in the pull-down are shown in the right hand panel. D, IL-1β levels in the supernatants 6 h post infection with indicated *T. gondii* strains was determined in wt and *Gbp5* ko BMDCs in the presence of IFN-γ. IL-1β levels are significantly decreased in ko cells compared to wt cells upon *T. gondii* wt strain infection. E, Nigericin treatment served as positive control for inflammasome activation. No significant differences can be observed in wt compared with ko cells. F, No differences in LDH release are apparent in wt compared with ko cells upon infection with *T. gondii* wt or ko strains. D, E, F, Error bars indicate the mean and standard error of the mean (SEM) of five or seven independent experiments. D, Multiple unpaired Student’s t-test was used for two-group comparisons; *p < 0.05.

Identification of ROP5 and ROP18 as inhibitors of inflammasome activation, along with the association of ROP5 with GBP5, implicates contribution of GBP5 to inflammasome activation upon *T. gondii* infection. A direct, potentially inhibitory association with ROP5 could be responsible for the differences in IL-1β secretion between *T. gondii* wt and ko strains observed in CIM-derived BMDCs (Fig. 5). Wt and *Gbp5* ko BMDCs (Suppl. Fig. 4) were seeded in the presence of IFN-γ and LPS and infected for 6 h with *T. gondii* RHΔ*hxgprt*, RHΔ*rop5* or RHΔ*rop18*. IL-1β levels were determined in the supernatants by quantitative ELISA. Compared to wt cells, IL-1β levels are significantly reduced in *Gbp5* ko BMDCs upon infection with *T. gondii* wt but not with RHΔ*rop5* or RHΔ*rop18* strains (Fig. 6D). Nigericin served as positive control for inflammasome activation (Fig. 6E). These results suggest that ROP5/ROP18, via its interaction with GBP5, directly inhibits GBP5-dependent inflammasome activation. However, the level of cell death, determined by LDH release, is not affected by the absence of *Gbp5* (Fig. 6F).

### GBP5-dependent inflammasome activation in human macrophages is not inhibited by *T. gondii* ROP5 and ROP18

To investigate the contribution of GBP5 to inflammasome activation upon *T. gondii* infection of human macrophages, we generated THP-1 *Gbp5* ko cells by CRISPR/Cas9 (clustered regularly interspaced short palindromic repeats/CRISPR-associated protein 9) technology. The absence of GBP5 was confirmed in detergent lysates after IFN-γ-stimulation for 24 h with a GBP5-specific antibody in comparison with wt cells. Expression levels of GBP3 served as specificity control and were essentially the same in all conditions (Fig. 7A).

**Figure 7.**
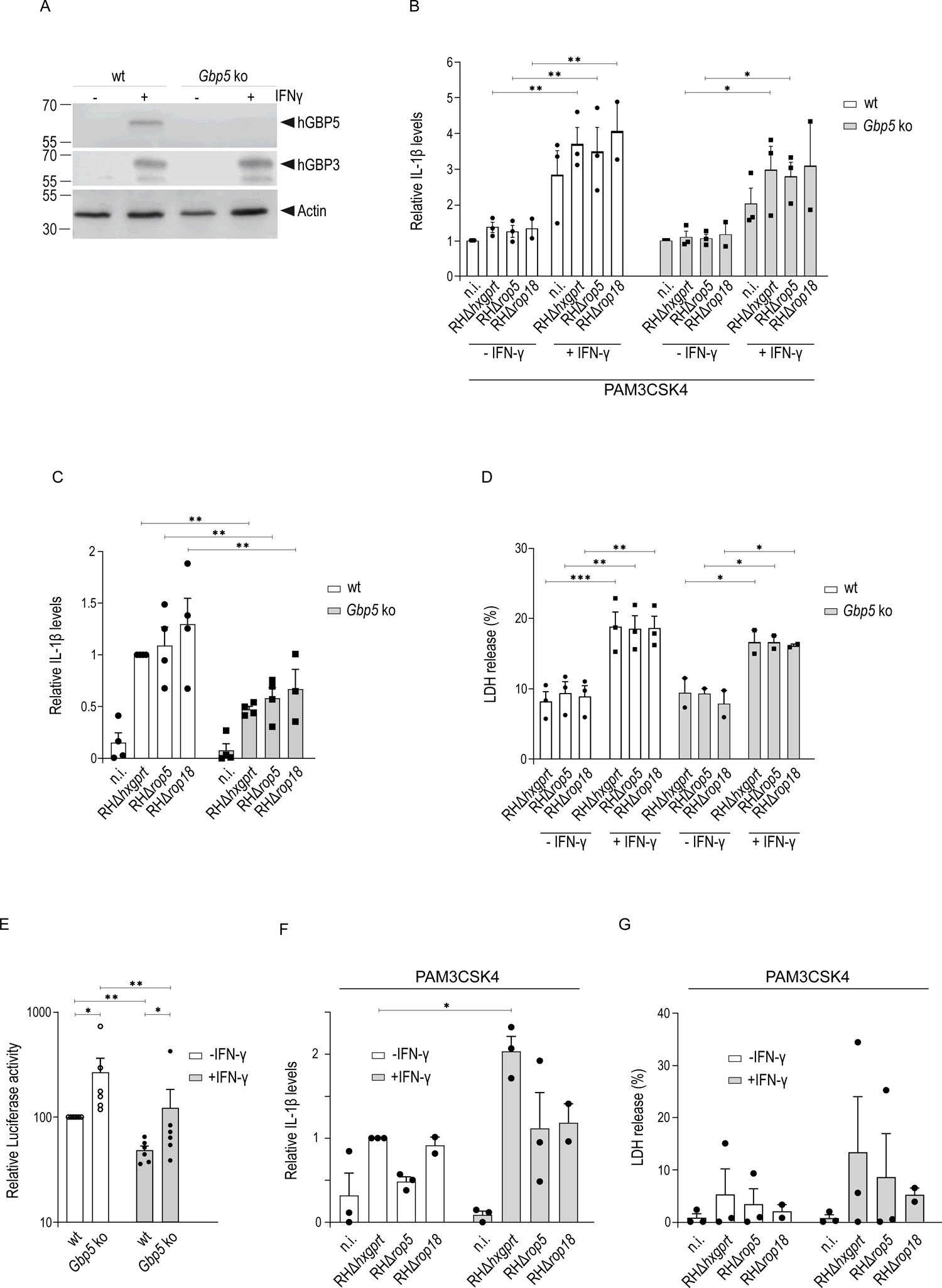
GBP5-dependent inflammasome activation in human macrophages is not inhibited by *T. gondii* ROP5 and ROP18. A, Western blot of detergent lysates from THP-1 wt and *Gbp5* ko cells stimulated for 24 h with IFN-γ. Compared with wt cells, hGBP5 expression is completely absent in ko cells (upper panel). GBP3 expression is unchanged in wt compared with ko cells (middle panel). Actin serves as loading control (lower panel). B, C, D, THP-1 macrophages were prestimulated for 3 h with LPS (B) or not (C, D) and subsequently infected with indicated *T. gondii* strains in presence or absence of IFN-γ for 6 h. No differences in IL-1β levels in the supernatant are apparent between *T. gondii* wt, RHΔ*rop5* and RHΔ*rop18* infections (B, C). Without LPS priming, IL-1β secretion is significantly decreased in the absence of GBP5 compared with wt cells upon infection with all *T. gondii* strains. D, Cell death, determined by LDH release into the supernatant, is significantly increased in the presence of IFN-γ compared with unstimulated conditions upon infection with all *T. gondii* strains but not differences between wt and *Gbp5* ko cells are detectable. E, THP-1 macrophages were stimulated with IFN-γ for 24 h or left unstimulated and infected with *T. gondii* RH expessing firefly luciferase (ME49-GFP-Luc) for 48 h. Parasite replication, determined as described in Material and Methods, is significantly increased in the absence of GBP5 but significantly reduced in the presence of IFN-γ compared with unstimulated conditions. F, G, Macrophages prepared from three healthy human donors were prestimulated for 3 h with LPS and subsequently infected with indicated *T. gondii* strains in presence or absence of IFN-γ for 6 h. IL-1β levels in the supernatants are significantly increased in case of *T. gondii* RHΔ*hxgprt* upon IFN-γ stimulation compared with unstimulated conditions (F). No differences in cell death, determined by LDH release into the supernatant, are apparent between *T. gondii* strains in the absence or presence of IFN-γ (G). B, C, D, E, F, G, Error bars indicate the mean and standard error of the mean (SEM) of two, three, four or six independent experiments. One-way analysis of variance (ANOVA) followed by Tukey’s (B, G) or Sidak’s (E) multiple comparison was used to test differences between groups; **p < 0.01, *p < 0.05.

Wt and *Gbp5* ko THP-1 macrophages seeded in the absence or presence of IFN-γ and after prestimulation with the TLR ligand PAM3CSK4 for 3 h were infected with *T. gondii* RHΔ*hxgprt*, RHΔ*rop5* or RHΔ*rop18* for 6 h and IL-1β levels determined by ELISA. IL-1β levels in the supernatants of *T. gondii* wt and ko strain-infected cells remain the same regardless of the presence of IFN-γ and no differences between *T. gondii* strains are apparent (Fig. 7B). Because of high IL-1β levels in the SN of even uninfected cells, which could mask differences between *T. gondii* strains, we performed a repeat experiment in the presence of IFN-γ but without prior PAM3CSK4 prestimulation. In this setup, almost no IL-1β release is observed in non-infected conditions but differences between *T. gondii* strains are still not detectable (Fig. 7C). However, infection of *Gbp5* ko cells with all *T. gondii* strains induces less IL-1β secretion than in wt cells (Fig. 7C) demonstrating the importance of GBP5 for inflammasome activation upon *T. gondii* infection of human macrophages. While cell death, determined by LDH release, is increased in the presence of IFN-γ, any variations between wt and ko *T. gondii* strains are not apparent (Fig. 7D). When we reinvestigated replication of *T*. *gondii* type II expressing firefly luciferase (ME49-GFP-Luc) in THP-1 macrophages [35], inhibition of replication is significantly diminished in *Gbp5* ko relative to wt cells in presence and absence of IFN-γ (Fig. 7E). To determine a possible inhibitory effect of ROP5/ROP18 on inflammasome inhibition in a different *in vitro* model system, we repeated the experiment in primary human macrophages of three different healthy blood donors. Under these conditions, even after prestimulation with PAM3CSK4 for 3 h, almost no IL-1β in the SN is detectable in non-infected cells. In infected cells, IL-1β release but not cell death is significantly increased upon IFN-γ stimulation compared with unstimulated conditions but just like in THP-1 cells, there was no effect of ROP5/ROP18 deficiency on IL-1β secretion (Fig. 7F) or cell death (Fig. 7G) is visible. These results emphasize a role for GBP5 in inflammasome activation during *T. gondii* infection, while at the same time stressing that the *T. gondii* ROP5/ROP18 virulence system is probably highly significant for the interaction between the parasite and murine rodent species.

## Discussion

Inflammasome activation is important in controlling microbial infections and many pathogens modulate inflammasomes to promote immune evasion and to facilitate dissemination. Whereas NLRP1 and AIM2 inflammasomes are activated by direct interaction with specific pathogen associated molecular patterns (PAMPS), the NLRP3 inflammasome is activated by a large number of chemically and structurally diverse agonists [59, 60]. Bacteria [61], fungi [62], viruses [63] and parasites [64] all have been demonstrated to activate the NLRP3 inflammasome. It is therefore unlikely that this sensor directly detects specific stimuli but rather responds to cellular stress that is induced by these infectious agents.

It has been demonstrated for different vacuolar bacteria that inflammasome-mediated immunity is promoted by Interferon (IFN)-γ-inducible products [8, 11, 12, 65–67]. We here demonstrate that priming with IFN-γ increases interleukin (IL)-1β secretion and cell death also in *T. gondii* infected bone marrow-derived dendritic cells (BMDCs). This observation contradicts other findings [47] and could be due to the fact that in the study by Ewald et *al*. any possible contribution of IFN-γ was investigated without NLRP3 licensing (TLR ligands). Accumulation of IFN-γ-inducible Immunity-Related GTPases (IRG proteins) and Guanylate Binding Proteins (GBP proteins) at the parasitophorous vacuolar membrane (PVM) of *T. gondii* in mice eventually leads to membrane disruption and parasite death. Without IRG protein loading, vacuolar disruption, unvariably followed by parasite and host cell death, has never been observed [29]. Effector IRG proteins that accumulate at the PVM are regulated by IRG proteins Irgm1, Irgm2 and Irgm3 which keep the effector IRG proteins in an inactive GDP-bound state in the host cell cytosol or at endomembranes in uninfected cells. In the absence of IRGM proteins, IRG and GBP proteins do not accumulate at the PVM and membrane integrity is therefore preserved [50–54]. In *Irgm1/Irgm3* ko BMDCs, percentages of cell death and IL-1β levels were significantly reduced compared to wt cells and differences for individual *T. gondii* strains were not apparent anymore. These results confirm the regulatory defect of the IRG/GBP system in the absence of Irgm1/Irgm3 resulting in preservation of vacuolar membrane integrity and suggest that IRG/GBP protein-mediated PVM disruption is responsible for NLRP3 inflammasome activation upon infection. Low levels of IL-1β secretion were detected even in *Irgm1*/*Irgm3* ko cells. An explanation would be residual levels of PVM disruption even in the absence of Irgm1/Irgm3 that is also reflected by residual cell death under these conditions (Fig. 3). Triple Irgm1/Irgm2/Irgm3-deficient cells [53, 54] should reveal the overall impact of IRG- and GBP-mediated PVM destruction on inflammasome activation upon *T. gondii* infection. It is known that pathogen challenge can lead to activation of multiple inflammasomes [68–70] and residual IL-1β release in our experiments in the absence of NLRP3 would confirm activation of the NLRP1 [36, 37] and/or AIM2 inflammasome [49] by *T. gondii*. However, a large part of the inflammasome response upon infection of BMDCs in our experiments is due to NLRP3 activation downstream of vacuolar disruption.

Differences of gasdermin-D (GSDMD) cleavage, an oligomeric pore-forming protein mediating pyroptosis upon proteolytic cleavage of a flexible linker between a cytotoxic N-terminal and a C-terminal repressor domain [71], upon wt and ko strain infections do not result in different levels of cell death in our experiments. These results are consistent with lytic but non-pyroptotic cell death upon *T. gondii* infection [29, 37, 47] that is mediated by IRG/GBP accumulation at the PVM and allows IL-1β secretion. *T. gondii*-mediated NLRP3 activation and release of IL-1β without pyroptosis has also been demonstrated in human monocytes [72, 73]. However, we cannot entirely dismiss the possibility that subtle levels of pyroptosis may be present but masked by this alternative form of lytic cell death.

The molecular mechanisms that allow maintenance of *T. gondii* PVM integrity are fairly well understood. Virulence of *T. gondii* is largely determined by allelic differences in secreted effector proteins that protect the PVM from IRG/GBP-mediated disruption. ROP5 scaffolds other parasite effectors - e.g. ROP18 - in multiprotein complexes to achieve efficient IRG protein inactivation. Protection of PVM integrity by *rop5*/*rop18* type I alleles explains diminished IL-1 β levels in wt compared with *T. gondii rop5* or *rop18* ko infections and confirms PVM disruption as important trigger for inflammasome activation. In some wild-derived Eurasian mice (such as the *Mus musculus castaneus* substrain CIM), the polymorphic tandem IRG protein Irgb2-b1 directly binds to ROP5 isoform B (ROP5B). Consequently, the PVM is vulnerable to IRG/GBP-mediated disruption and replication even of *T. gondii* type I strains is efficiently inhibited [57, 58]. To provide additional evidence that NLRP3 inflammasome activation upon *T. gondii* infection is dependent on PVM disruption, we infected CIM-derived BMDCs and determined lactate dehydrogenase (LDH) and IL-1β release into the supernatant. Consistent with CIM-intrinsic resistance towards *T. gondii* type I strains, no differences in cell death were observed between *T. gondii* strains. However, ROP5/ROP18 significantly inhibited IL-1β release even under these conditions. For proper assembly of NLRP3 inflammasomes, homotypic interactions are required [74]. Enhanced ASC oligomerisation in the absence of either ROP5 or ROP18 in CIM cells indicates that the molecular mechanism of inflammasome inhibition by ROP5/ROP18 is inhibition of inflammasome complex formation rather than acting as an antagonist of caspase-1 activity, a strategy employed by viruses, bacteria [75] and fungi [76]. To conclude, *T. gondii* ROP5/ROP18 promote parasite survival by protection of the intracellular replicative niche, the PV, and by actively subverting NLRP3 activation. These findings are concomitant with a dual role of ROP5 and ROP18 for inflammasome inhibition in murine *T. gondii* infections.

The interaction between GBP5 and NLRP3 is crucial for NLRP3 complex assembly in human and mouse macrophages in response to pathogenic bacteria *Salmonella typhimurium* and *Listeria monocytogenes* or soluble priming agents [8]. We could here demonstrate reduced IL-1β secretion from *Gbp5* ko BMDCs demonstrating a critical role of GBP5 for efficient inflammasome activation even upon *T. gondii* wt strain infection. No differences in cytokine levels were apparent upon *T. gondii* ko strain infections of *Gbp5* ko compared with wt cells, probably because inflammasome activation triggered by PVM disruption in these cases masks any GBP5-releated effect. No differences in terms of cell death between wt and *Gbp5* ko BMDCs could be observed indicating that GBP5 does not significantly contribute to vacuolar disruption resulting in parasite and host cell death. This is in agreement with previous studies reporting that only 3-5 % *T. gondii* type II- and type III-derived vacuoles were found to be positive for GBP5 [6, 77]. No interaction between a *T. gondii* effector and a GBP protein has been identified yet. Here we show direct interaction of *T. gondii* ROP5A and ROP5B with hGBP5 and mGBP5. This Interaction suggests that direct inhibition of inflammasome complex formation by ROP5A/B (Fig. 5) is mediated by preventing the molecular function of mGBP5. It remains to be determined whether ROP5 binding to mGBP5 covers the interface crucial for NLRP3 and/or ASC interaction or - not mutually exclusive - enables recruitment of the active kinase ROP18 that inhibits inflammasome activation via phosphorylation of GBP5 or any other complex component. We have demonstrated earlier that ROP5B is the most relevant isoform that determines parasite virulence in mice [57]. Here we show interaction of ROP5A and ROP5B with mGBP5. It is currently unclear which of these ROP5 isoforms, if not both, are required for NLRP3 inhibition upon *T. gondii* infection.

Since almost complete loss of the IRG resistance system [78], humans rely on other effector mechanisms against *T. gondii*. In THP-1 cells, GBP1, GBP2 and GBP5 have been demonstrated to control *T. gondii* infection [35]. Whereas GBP1 accumulation at the *T. gondii* type I and type II strain-derived PVM is required for vacuolar disruption and lysis of the parasite plasma membrane in different macrophages, GBP2 and GBP5 perform a pathogen-distal role in parasite restriction in these cells. In contrast, about 60 % GBP5-positive type II strain-derived PVs have been found in HeLa cells [79]. Future experiments should reveal the impact of GBP5 on vacuolar disruption in different human cell types and whether PVM disruption - in analogy to BMDCs - triggers inflammasome activation. We confirm the role of hGBP5 for *T. gondii* control [35] in our experiments (Fig. 7G). Furthermore, we show that in the absence of hGBP5, inflammasome activation and IL-1β release is significantly inhibited. Whether this finding reflects the importance of hGBP5 for *T. gondii* control or whether hGBP5 encompasses additional roles that extend beyond the scope of this observation can not be conclusively answered yet. In contrast to results observed in BMDCs and consistent with previous published results [42], ROP5 and ROP18 do not inhibit IL-1β secretion in human macrophages. However, the direct interaction with ROP5A/B (Fig. 6) could imply that hGBP5 is possibly inhibited by one or both of these parasite effectors, irrespective of its possible additional function in controlling *T. gondii* infection.

Regulation of inflammasome activation is crucial for controlling infection and preventing disease progression. Our work provides further insight into inflammasome activation and inhibition upon *T. gondii* infection. A more comprehensive understanding of the molecular details that contribute towards regulation of inflammasome activation could aid in the development of new strategies to improve clinical outcomes.

## Propagation of *T. gondii* strains

Tachyzoites of *T. gondii* strains RHΔ*hxgprt*, RHΔ*rop18*, RHΔ*rop5* and ME49-GFP-Luc [1] were maintained by serial passage in confluent monolayers of human foreskin fibroblasts (HS27, ATCC CRL-1634). After differential centrifugation, tachyzoite pellets were filtered through a 5 µm filter and washed two times in complete RPMI medium (Gibco) supplemented with 100 U/ml penicillin, 100 mg/ml streptomycin (Gibco) and 10 % fetal bovine serum (Anprotec). For infection experiments, purified tachyzoites were resuspended in complete RPMI medium or complete RPMI medium containing 20 ng/mL GM-CSF (Immunotools) respectively. The filtered parasites were immediately used for infection of cells or lysed for subsequent pull-down experiments.

## Cell culture

HEK293T cells and human foreskin fibroblast (HS27, ATCC CRL-1634) were maintained by serial passage in DMEM, high glucose (Invitrogen Life Technologies) supplemented with 100 U/ml penicillin, 100 mg/ml streptomycin (PAN) and 10 % fetal bovine serum (Anprotec). THP-1 cells were maintained in complete RPMI medium.

## Preparation and differentiation of mouse bone marrow cells, human PBMC-derived macrophages and THP-1 cells

Bone marrow cells were flushed from tibias and femurs with a 26 G needle, filtered through a 100 µm cell strainer and pelleted for 5 min at 450 g. To remove erythrocytes, cell pellets were resuspended in 1 mL of RBC lysis buffer (Qiagen) and incubated for 5 min at RT. After addition of 9 mL of flush-medium, cell pellets (450 g, 5 min, RT) were resuspended in 10 mL DC-medium (20 ng/ml GM-CSF) at 5 x 10^5^ cells/ml. 10 ml of the cell suspension were differentiated to BMDCs (Bone Marrow Derived Dendritic Cells) for 7 days in a 10 cm petri dish containing DC medium. At day 2 and 4 of differentiation, the medium was replaced with 10 ml of fresh DC-medium.

2.5 x 10^5^ THP-1 cells were differentiated in 24 well plates to macrophages with 100 ng/mL PMA o/n. Cells were washed with DPBS (Gibco) and medium replaced before infection. For PAM3CSK4 priming, THP-1 cells were differentiated for 24 h with 100 ng/mL PMA. After a washing step with DPBS, THP-1 cells were incubated with RPMI complete medium for two additional days before infection.

PBMCs were isolated by gradient centrifugation using Ficoll-paque PLUS (Cytiva) from healthy donors. 2 x 10^6^ cells were seeded in 24 well plates in complete RPMI medium supplemented with 40 ng/mL M-CSF (PeproTech). 24 h after seeding, non-adherent cells were removed by two washing steps with PBS and fresh medium containing M-CSF was added to each well every two days for 6 days.

## Stimulation, priming and infection of BMDCs, PBMC-derived macrophages and THP-1 cells

24 h before infection, differentiated BMDCs were harvested with HBSS/EDTA for 10 min at 37°C and 5 % CO_2_ and cell pellets resuspended in fresh DC-medium. BMDCs (0.5 x 10^4^ to 1 x 10^5^ cells in each well of a 96 well plate), PBMC-derived macrophages and THP-1 cells were stimulated (or left unstimulated) with 100 ng/ml IFN-α or 100 U/mL IFN-γ for 24 h and primed with LPS (20 ng/mL) (Sigma) or PAM3CSK4 (100 ng/mL) (Invivogen) (or left unprimed) for 3 h before infection with different *T. gondii* strains for 6 h (BMDCs and PBMC-derived macrophages) or 24 h (THP-1 cells). Nigericin (5 µM) (Sigma) was added as a positive control for inflammasome activation 0.5 - 1 h before cells were harvested by centrifugation for 5 min at 400 g. Supernatants and pellets were used immediately for ELISA, LDH release assay or Western blot or stored at −20°C.

## LDH release assay

To quantify cell death, the Pierce LDH Cytotoxicity Assay Kit (Thermo Fisher Scientific) was used. 50 µL of supernatants were transferred into a well of a 96 well plate and incubated with 50 µL of the reaction mixture for 30 min at RT. The reaction was stopped by adding 50 µL stop solution, the absorbance measured at 490 nm and corrected by subtracting the absorbance at 680 nm. 1x lysis buffer was used to prepare a positive control for cell death. The percentage of cell death was calculated using the formula: 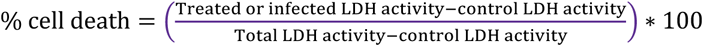

## IL-1β sandwich ELISA

IL-1β in the supernatants was determined with commercial sandwich ELISA kits. For BMDCs, the mouse IL-1β/IL-1F2 DuoSet ELISA kit (R&D Systems) was used. For PBMC-derived macrophages and THP-1 cells, the ELISA MAX™ Deluxe Set Human IL-1β (BioLegend) was used. TNF-α was determined using the ELISA MAX™ Standard Set Mouse TNF-α kit (BioLegend). The absorbance was determined at 450 and 570 nm and the concentration of cytokines calculated from a standard curve.

## ASC cross-linking and oligomerisation assay

2 x 10^6^ BMDCs were seeded in a 6 well plate, stimulated or not with 100 U/mL IFN-γ and primed with LPS (20 ng/mL) for 3 h before infection with different *T. gondii* strains for 2 h. Nigericin (5 µM) was added as a positive control for inflammasome activation for 0.5 - 1 h. Cells were washed 1 x with ice-cold PBS before resuspension in 0.8 mL of ice-cold lysis buffer (20 mM HEPES, 10 mM KCL, 1mM MgCl_2_, 1% NP-40, 1 mM DTT, Benzonase 25 U/mL, 1x Complete protease inhibitor cocktail (Roche)). Cells were lysed with a 21-gauge needle (10x) and incubated for 15 min at 4°C under continuous rotation. A 40 µl aliquot of the lysate was removed and used for subsequent Western blot analysis. Cell lysates were centrifuged at 3,000 g for 10 min at 4°C before the pellets were finally resupended in ice-cold PBS containing 2 mM DSS (Thermo Fisher Scientific). After 30 min at RT and addition of 1 ml ice-cold PBS, pellets (16000 g, 10 min, 4°C) were resuspended in 40 µl 1x Laemmli sample buffer and ASC oligomerization was analysed using an ASC-specific antibody by Western blot.

## Expression and purification of recombinant GST-hGBP5

Recombinant GST-fusion protein hGBP5 was expressed and purified as described previously [2].

## Yeast Two-Hybrid assay

Saccharomyces cerevisiae strain PJ69-4α was incubated with 1 µg of plasmid DNA (pGAD-C3 or pGBD-C3 containing the indicated genes) in transformation buffer (40 % PEG 3350, 0.2 M LiAc, 0.5 mg/ml ss DNA, 0.1 M DTT) for 30 min at 42°C. Cotransformants were selected by plating on double dropout media (SD/-Leu/-Trp). Colonies grown on double dropout media were replica plated again on double dropout media before OD_600_ measurement of single colonies resuspended in liquid triple dropout media (SD/-Leu/-Trp/-His). An equal amount of material was plated on triple dropout media containing 1 mM 3-AT and incubated for 5 to 10 days at 30°C.

## Plasmid Constructs

The pGEX-4T-2-hGBP5 construct allowing expression of recombinant GST-hGBP5 was generated after amplification of hGBP5 from cDNA prepared from HELA cells and ligation into pGEX-4T-2. hGBP5 and mGBP5 were amplified from pGEX-4T-2-hGBP5 and pWPXL-mGBP5 [3] respectively, and ligated into pGAD-C3. The pGBD-ROP5A/B/C constructs were generated earlier [4].

hGBP5 ko THP-1 cells were generated with the LentiCRISPRv2 (Addgene) plasmid CRISPR/Cas9 system. The insert fragment of *Gbp5* gRNA was generated by annealing the oligonucleotides hGBP5 1 (5′-caccgGAACTTTAATGAGCAGCTGA) and hGBP5 2 (5′-cTCAGCTGCTCATTAAAGTTCcaaa) and cloning into the Esp3I (BsmBI) (Thermo Fisher Scientific) site to the generate the gRNA-expressing plasmid LentiCRISPRv2_hGBP5.

## Lentiviral trasduction

Lentiviral transduction was applied to generate hGBP5 ko THP-1 cells. For this purpose, gag-pol-expressing and env-expressing plasmids were co-transfected with the plasmid carrying the gRNA (LentiCRISPRv2_hGBP5) [5] into HEK293T cells that have been grown to a density of 70 % in a 10 cm plate. 24 h post transfection, the medium was exchanged, and cells incubated for additional 24 h. The supernatant was filtered and transferred to 2,5 x 10^5^ THP-1 cells that have been seeded one day before in a 12 well plate. After 24 h, cells were harvested and transferred into appropriate cell culture flasks with medium containing 1 µg/ml puromycin for selection of transduced cells. After 1 week of selection, 1 cell/100µl was seeded into a well of a 96-well plate and each well observed for several days to contain only one single colony. Single colonies were transferred to 24 well plates and detergent lysates of IFN-γ-induced (100 U/ml) cells analyzed 24 h later by Western blot for hGBP5 expression.

## Immunological Reagents

Immunoreagents used in this study were: 3E2 mouse anti-ROP5A/B [6], 2.4.21 rat anti-*T. gondii* GRA7 [4], 10E7 mouse anti-Irga6 [7], B34 mouse anti-Irgb6 [8], goat anti-GST (GE Healthcare), goat anti-IL-1β (IL-1β/IL-1F2) (R&D Systems), mouse anti-caspase-1 (p20) (AdipoGen Life Sciences), rabbit anti-ASC (AL177) (AdipoGen Life Sciences), rabbit anti-GSDMD (abcam), mouse anti-human GBP5 (LS Bio), rabbit anti human-GBP3 (Abcam), rabbit anti-mouse GBP5 (Eurogentec), rabbit anti-calnexin (Merck) and mouse anti-actin (Sigma Aldrich). Alexa 488-labelled goat anti-rabbit and Alexa 555-labelled donkey anti-mouse fluorescent antisera (Life Technologies), goat anti-rabbit-HRP (Jackson Immuno Research Laboratories), goat anti-rat-HRP (Jackson Immuno Research Laboratories), donkey anti-goat-HRP (Abcam) and rabbit anti-mouse-HRP (Jackson Immuno Research Laboratories) polyclonal antibodies were used as secondary reagents.

## Postnuclear lysate preparation from free tachyzoites and infected cells

10-25 x 10^6^ *T. gondii* tachyzoites were washed trifold with PBS before lysis in 800 µl NP-40-lysis buffer (0.1 % NP-40, 150 mM NaCl, 20 mM Tris/HCl (pH 7.6), 5 mM MgCl_2_ supplemented with protease inhibitors (Roche)) for 2 h under constant rotation at 4°C. Postnuclear lysates were subjected to Western blot or pull-down analysis.

## Pull-Down Analysis

Purified GST (100 pmol) or GST-hGBP5 fusion protein (100 pmol) was incubated with 100 µl 1:1 (PBS/2 mM dithiothreitol (DTT)) glutathione sepharose 4B (GE Healthcare) resin suspension in a final volume of 600 µl PBS/2 mM DTT or 2 h at 4°C under constant rotation. The resin was washed trifold with ice-cold lysis buffer containing 2 mM DTT without detergent and incubated with 600 µl postnuclear tachyzoite lysate o/n at 4°C under constant rotation. Beads were washed trifold with ice-cold lysis buffer and either stored at −80°C or immediately boiled in sample buffer (80 mM Tris/HCl (pH 6,8), 5 mM EDTA, 4 % SDS, 34 % sucrose, 40 mM DTT, 0.002 % bromphenol blue) for 5 min at 95°C and subjected to SDS-PAGE and Western Blot.

## *T. gondii* replication assay

*T. gondii* replication was determined in THP-1 cells stimulated o/n or not with 100 U/mL IFN-γ (Peprotech) and infected with *T. gondii* ME49-GFP-Luc at a MOI of 0.25 for 48 h. Cells were washed once with PBS and lysed for at least 1 h with 40 µl 1x passive lysis buffer (Promega) at RT. 20 µl of lysate were transferred to a white flat-bottom 96-well plate (Thermo Fisher Scientific) and luciferase activity was measured in a Tecan infinite 200Pro by automatic injection of 50 µl luciferase assay substrate (Promega) and 10 sec integration time. Luciferase activity was expressed relative to non-stimulated wt THP-1 cells.

## Immunocytochemistry

Fixation and staining of BMDCs grown on coverslips was performed as described earlier [9]. Microscopy and image analysis was performed blind on coded slides essentially according to [10]. Intracellular parasites were identified from the pattern of *T. gondii* GRA7 staining.

## Statistics

All statistical analyses were performed using GraphPad Prism 9.1 software. P-values were determined by an appropriate statistical test. One-way ANOVA followed by Tukey’s, Sidak’s or Holm-Sidak’s multiple comparison was used to test differences between three or more groups. Depending on the data distribution, Multiple unpaired Student’s t-test was used for two-group comparisons. All error bars indicate the mean and standard error of the mean (SEM) of at least three independent experiments. P-values; ****p < 0.0001, ***p < 0.001, **p < 0.0, *p < 0.05, n.s. no significant.

## Ethics Statement

All animal experiments were performed in compliance with the German animal protection law (TierSchG). Mice were handled in accordance with good animal practice as defined by FELASA and the national animal welfare body GV-SOLAS. The animal welfare committees of the University of Freiburg as well as the local authorities (Regierungspräsidium Freiburg; Landesamt für Natur, Umwelt und Verbraucherschutz Nordrhein Westfalen; Behörde für Soziales, Familie, Gesundheit und Verbraucherschutz, Hamburg) approved all animal experiments.

## Data availability

The authors declare that all data supporting the findings of this study are available within the article and its Supplementary Information files, or are available from the authors upon request.

## Acknowledgments

We would like to thank Julia Mock very much for the preparation and provision of examination materials. Georg Häcker and Ian Gentle generously provided materials and technical assistance. Kerstin Flämig provided technical assistance with protein purification. We are thankful for all support from the Institute of Virology under the direction of Hartmut Hengel. T.S. received funding from the DFG (STE 2348/2) and the Research commission of the Faculty of Medicine Freiburg (STE 1113/17). O.Gr. received funding from SFB 1160 (Project ID 256073931), SFB/TRR 167 (Project ID 259373024), SFB 1425 (Project ID 422681845), SFB 1479 (Project ID 441891347), GRK 2606 (Project ID 423813989), as well as by the European Research Council (ERC) through Starting Grant 337689, Proof-of-Concept Grant 966687, the EU-H2020-MSCA-COFUND EURIdoc programme (No. 101034170) and from Germany’s Excellence Strategy, CIBSS – EXC-2189 (project ID 390939984). M.M.L. and S.S. received funding (Research Grants - Doctoral Programmes in Germany) from the German Academic Exchange Service (DAAD). The funders had no role in study design, data collection and interpretation, or the decision to submit the work for publication.

## Author Contributions

M.M.L. and T.S. conceived the study, M.M.L., A.M.B.Q., S.S., X.M., O.G., B.S., A.F.A.S. and T.S. designed the experiments, M.M.L., A.M.B.Q., S.S., X.M. B.S., A.F.A.S. and T.S. performed the experiments, M.M.L., A.M.B.Q. S.S., X.M., O.G., B.S., A.F.A.S., C.C., J.E.G.M, O.Gr., and T.S. evaluated the data. O. Gr., K.P., J.C.H. and G.A.T. provided reagents and examination materials. M.M.L., A.M.B.Q. and T.S. wrote the manuscript.

## Competing interests

The authors declare no competing interests.Tachyzoites of *T. gondii* strains RHΔ*hxgprt*

## Notes

### Competing Interest Statement

The authors have declared no competing interest.

