## Supplementary figures and images for "Dual role of *Toxoplasma gondii* ROP5 and ROP18 for NLRP3 inhibition"

### Suppl. Figure 1

**Suppl. Figure 1**

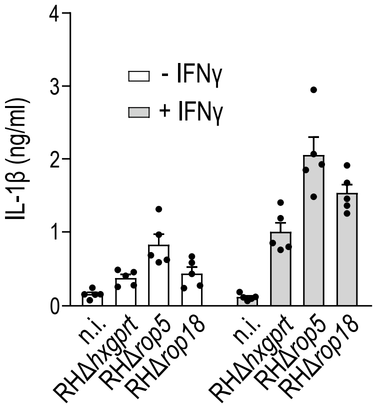

### Suppl. Figure 2

**Suppl. Figure 2**

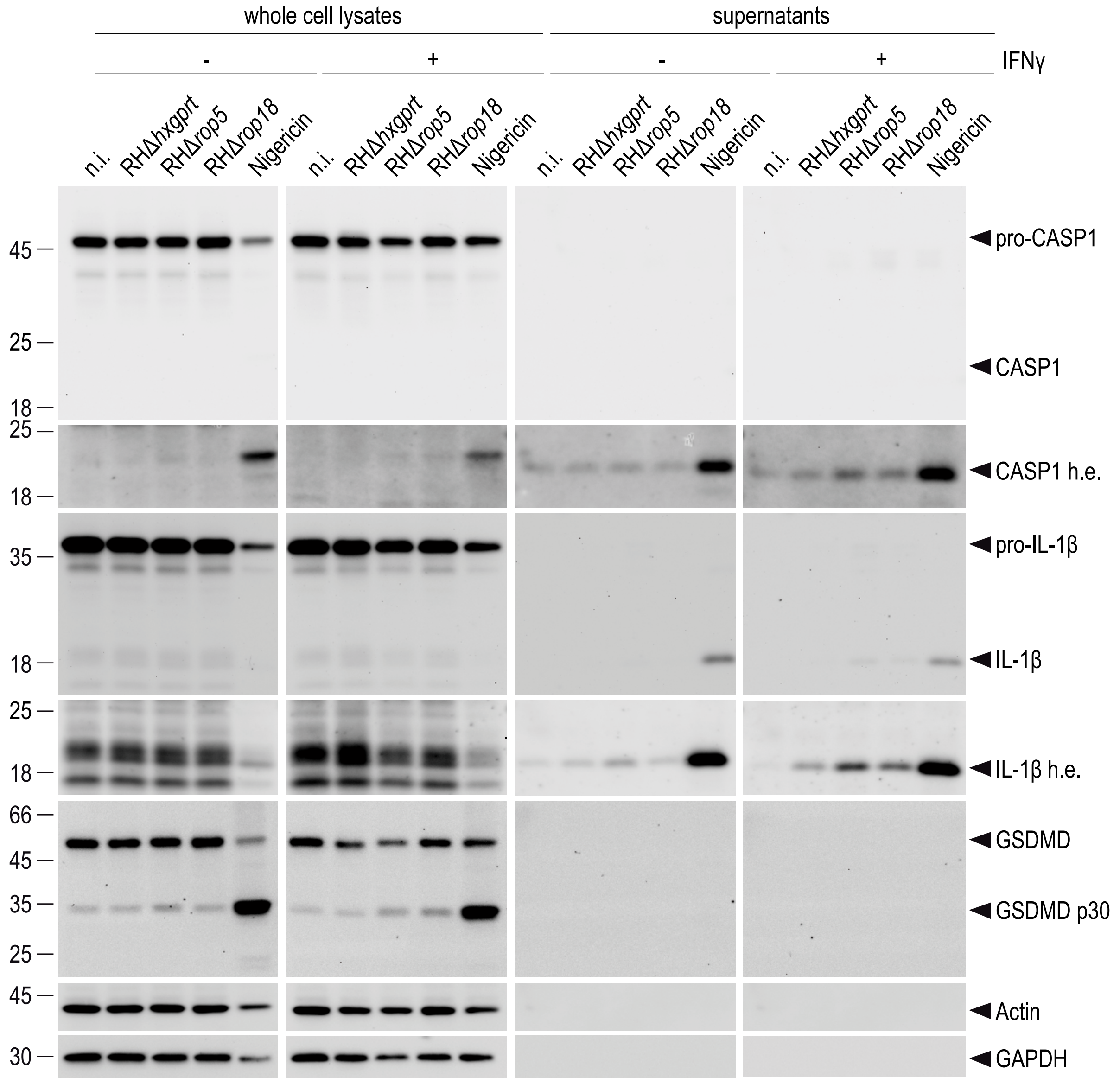

### Suppl. Figure 3

**Suppl. Figure 3**

A

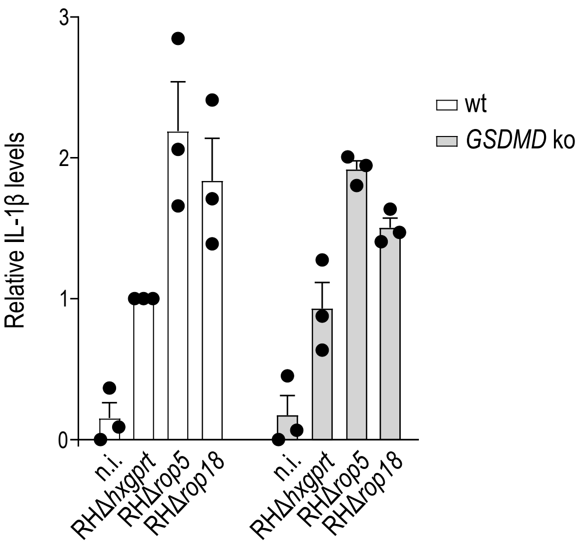


B

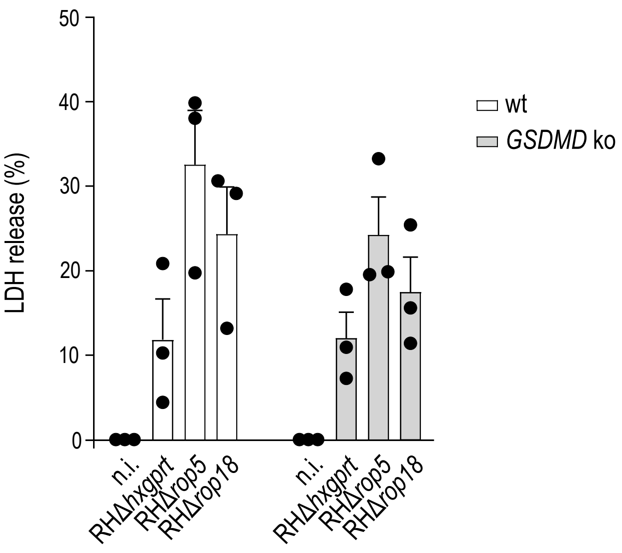

### Suppl. Figure 4

**Suppl. Figure 4**

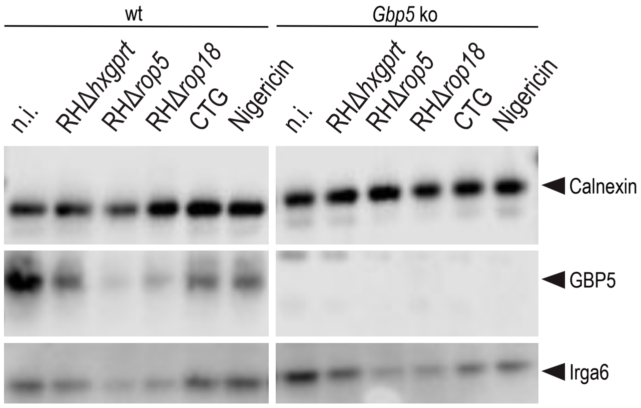
