## Supplementary material for "Dual role of *Toxoplasma gondii* ROP5 and ROP18 for NLRP3 inhibition": Suppl. Figure legends

**Suppl. Fig. 1. *T. gondii* ROP5 and ROP18 inhibit IL-1β production.** BL/6 BMDCs were primed for 3 h with LPS and subsequently infected with indicated *T. gondii* strains in presence or absence of IFN-γ for 6 h. IL-1β levels in the supernatants are significantly increased upon *T. gondii* infection in the presence of IFN-γ compared with unstimulated conditions. Comparing *T. gondii* strains, IL-1β secretion is significantly increased in ko compared to wt strain infections in the presence of IFN-γ. Non-infected cells served as negative control for inflammasome activation.

**Suppl. Fig. 2. IL-1β and gasdermin-D release into the supernatant is increased upon *T. gondii* RHΔ*rop5* and RHΔ*rop18* infection.** BL/6 BMDCs were primed with LPS and stimulated with IFN-γ. Cleavage of immature IL-1β (pro-IL-1β), caspase-1 (pro-CASP1) and gasdermin-D (GSDMD) and secretion of mature IL-1β (IL-1β, h.e.: higher exposure) and cleaved caspase-1 (CASP1, h.e.: higher exposure) and gasdermin-D (GSDMD p30) into the supernatant were analysed by Western Blot 6 h p.i. with indicated *T. gondii* strains.

**Suppl. Fig. 3. IL-1β and LDH release into the supernatant upon *T. gondii* infection is not mediated by gasdermin-D.** BL/6 wt or *GSDMD* ko BMDCs were primed with LPS and subsequently infected with indicated *T. gondii* strains in presence or absence of IFN-γ for 6 h. IL-1β (A) and LDH (B) release into the supernatant upon *T. gondii* infection is comparable in wt and ko cells.

**Suppl. Fig. 4. Analysis of wt and *Gbp5* ko BMDCs.** Western blot of detergent lysates from wt and *Gbp5* ko BMDCs stimulated for 24 h with IFN-γ and subsequently infected with indicated *T. gondii* strains for 2 h. Compared with wt cells, GBP5 expression is completely absent in ko cells (middle panel). Irga6 expression is unchanged in wt compared with *Gbp5* ko cells (lower panel). Calnexin serves as loading control (upper panel).
